# Intragenic differential expression in archaea transcriptomes revealed by computational analysis of tiling microarrays

**DOI:** 10.1101/163436

**Authors:** Atlas Khan, Ricardo Z. N. Vêncio

**Affiliations:** Universidade de São Paulo, LabPIB, Department of Computing and Mathematics, FFCLRP-USP, Av. Bandeirantes, 3900, Ribeirao Preto, SP 14040-901, Brazil; Universidade de São Paulo, LaBiSisMi, Department of Biochemistry and Immunology, FMRP-USP, Av. Bandeirantes, 3900, Ribeirao Preto, SP 14049-900, Brazil

**Keywords:** Archaea, tilling array, transcription, self-organizing map, gap statistics

## Abstract

Recent advances, in high-throughput technologies allows whole transcriptome analysis, providing a complete and panoramic view of intragenic differential expression in eukaryotes. However, intragenic differential expression in prokaryotes still mystery and incompletely understood. In this study, we investigated and collected the evidence for intragenic differential expression in several archaeal transcriptomes such as, *Halobacterium salinarum* NRC-1, *Pyrococcus furiosus, Methanococcus maripaludis*, and *Sulfolobus solfataricus*, based on computational methods; specifically, by well-known self-organizing map (SOM) for cluster analysis, which transforms high dimensional data into low dimensional. We found 104 (3.86%) of genes in Halobacterium salinarum NRC-1, 59 (2.56%) of genes in *Pyrococcus furiosus,* 43 (2.41%) of genes *Methanococcus maripaludis* and 13 (0.42%) of genes in *Sulfolobus solfataricus* have two or more clusters, i.e., showed the intragenic differential expression at different conditions.

## Introduction

Recently, huge amounts of data from high-throughput sequencing and tiling arrays have been used to annotated genome and produced novel transcripts [1, 2]. Microarrays are a progressive achievement in experimental molecular biology that can all the while study a large number of qualities under a huge number of conditions and give a mass of information to the scientist.

Intragenic differential expression is vital and play important rule in eukaryotes. Intragenic differential expression in eukaryotes exists due to splicing, overlapping, mis-annotation of genome [3-5]. However, in prokaryotes the rule of intragenic differential expression is still remains challenging and mysterious. Our group showed that there are overlapping sotRNAs [6, 7] are exist in archaea. Some of them are observed as differentially expressed in a single condition. Our hypothesis is that intragenic differential expression can be found in several experimental conditions and in other genes presenting overlapping RNAs not only sotRNA or TSSaRNA. We will call them generally alternative transcripts. There are lots of already publicly available dataset that could answer question above but that did not address the problem. Bioinformatics is the way to go.

Cluster analysis is strategy for recognizing homogeneous groups of objects called clusters, which resemble each other and which are different in some respects from individuals in other clusters [8, 9]. SOM is a kind of neural networks that trained by using unsupervised learning to produce low-dimensional of input n-dimensional space of training samples [10-12] and have a specific characteristic that make it well suited to clustering of gene expressions data over time at different experimental conditions [13]. One of the most important question in SOM that how to determine the suitable number of clusters in data, so we used Gap statistics [14] with SOM to estimate the number of patterns (clusters) presented in data. Since it is hard to collect intragenic differential expression candidates genes one by one, so we used the cluster technique based on SOM with Gap statistics to present the intragenic differential expression at different conditions in archaea.

Here we analyzed the tiling microarrays data to the present intragenic differential expression in archaea, such as to investigate (i) alternative transcript, (ii) mis-annotation of genome, (iii) overlapping transcripts in third domain of life archaea based on computational methods.

## Results and Discussion

In this section, we presented the main contributions of the paper. We discussed the intragenic differential expression in the third domain of life archaea by computational analysis of tilling microarrays. We investigated and collected the evidence for (i) alternative transcripts, (ii) miss-annotations, (iii) overlapping transcripts, in archaea by examining all publicly available gene expression data of archaea to date.

### Result 1

We re-analyzed all the publicly available data and visualize it in Geggle Genome Browser (GGB) [15].

### Result 2

We used GAP statistics and SOM to automatically select the candidates with intragenic differential expression in *Halobacterium salinarum* NRC-1. Since it is hard to see them one by one, so we used cluster technique to select the genes, which have more than one clusters, which is related to intragenic differential expression.

### Result 3

We manually collected all candidates in user friendly visual tool (Result 1) and mined putative cases of intragenic differential expression.

### Result 4

We predicted similar phenomena in other archaea i.e., *Pyrococcus furiosus, Methanococcus maripaludis*, and *Sulfolobus solfataricus*, which have less data but can show some cases.

### Discussion

Intragenic differential expression could be due to several things:

1. Alternative transcripts
2. Mis-annotations of genome
3. Overlapping of genes (sotRNAs and TSSaRNAs)
4. RNA degradation
5. Noise

We did filtering selection to select only those cluster patterns that make sense for example VNG1743C (Figure 1 (a)). In this gene, half part of it red (over-expressed) and other half part is green (lower-expressed). From it, we may conclude that it is differently expressed at different conditions. To avoid 5), we did eigen match analysis and separated the noisy genes Table 1. We presented and collected the evidence for the sense overlapping genes, which are the generalization of sotRNAs [6] and we tabulated them in Table 2 and (Figure 1 (b)). We also investigated and predicted the TSSaRNAs, which are the generalization of TSSaRNA [7] Table 3. and (Figure 1 (c)). Our analysis showed that there are some alternative transcripts exist in archaea, which we predicted and tabulated in Table 4 and (Figure 1 (d)). Our method also presents the mis-annotation of the arachea genome and we presented the mis-annotation of genes in Table 5.

**Figure 1:**
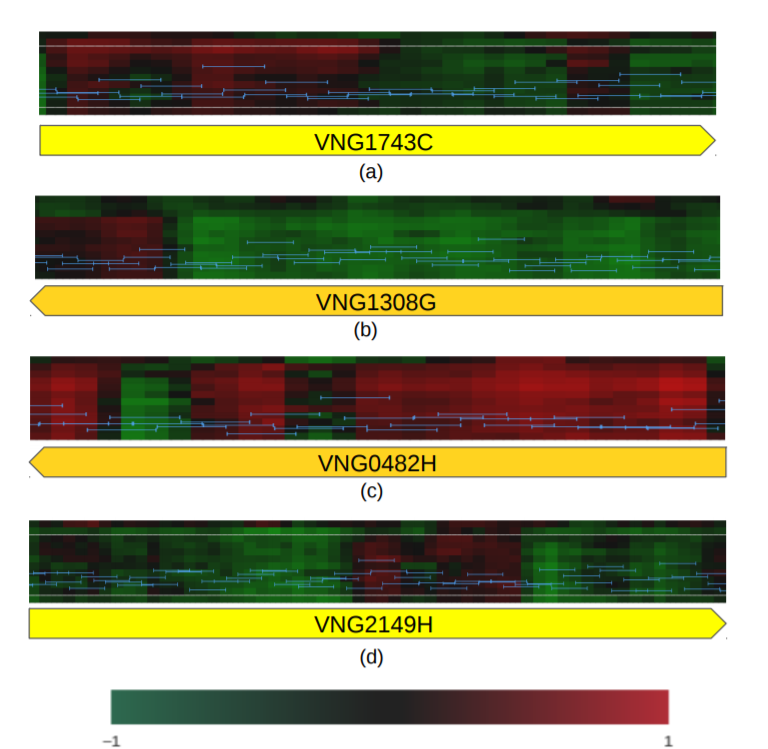
The intragenic differential expression of gene VNG1743C in H. salinarum NRC-1. The yellow arrow represents genes, horizontal axis represents organism’s genome coordinates, heatmap shows gene expression profiles over curves and color-coded according to ratios between each time point relative to reference condition. Light blue bar bars show tilling array probe intensities for experiment condition at time t(0). (a) shows the intragenic differential expression of gene VNG1743C, (b) shows sotRNAs, (c) shows TssaRNAs and (d) alternative transcripts of genes.

**Table 1:**
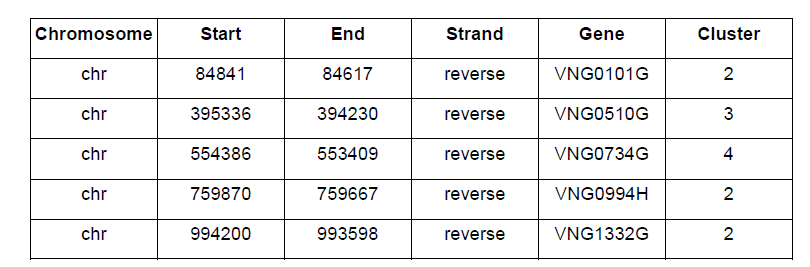
The list of noisy genes in *H. salinarum* NRC-1.

**Table 2:** The list of sotRNAs in *H. salinaru*m NRC-1.

| Chromosome | Start | End | Strand | Gene | Cluster |
| --- | --- | --- | --- | --- | --- |
| chr | 184345 | 184965 | forward | VNG0217H | 3 |
| chr | 871623 | 871844 | forward | VNG1151H | 2 |
| chr | 1065778 | 1066809 | forward | VNG1435G | 2 |
| chr | 1558353 | 1558970 | forward | VNG2121C | 3 |
| chr | 246854 | 246267 | reverse | VNG0314G | 2 |
| chr | 260251 | 258965 | reverse | VNG0329G | 3 |
| chr | 570320 | 569376 | reverse | VNG0752G | 3 |
| chr | 962319 | 962119 | reverse | VNG1283H | 2 |
| chr | 979291 | 978413 | reverse | VNG1308G | 2 |
| chr | 1333429 | 1332857 | reverse | VNG1798H | 2 |
| chr | 1448213 | 1447671 | reverse | VNG1962C | 2 |

**Table 3:** The list of TssaRNAs in *H. salinarum* NRC-1.

| Chromosome | Start | End | Strand | Gene | Cluster |
| --- | --- | --- | --- | --- | --- |
| chr | 1162079 | 1162777 | forward | VNG1559H | 3 |
| chr | 1698333 | 1699142 | forward | VNG2282C | 2 |
| chr | 45456 | 44245 | reverse | VNG0051G | 2 |
| chr | 117579 | 117313 | reverse | VNG0141H | 2 |
| chr | 373508 | 372939 | reverse | VNG0482H | 3 |
| chr | 374761 | 373577 | reverse | VNG0483C | 2 |
| chr | 1207039 | 1206584 | reverse | VNG1621H | 2 |
| chr | 1283685 | 1283350 | reverse | VNG1734H | 4 |
| chr | 2007438 | 2006920 | reverse | VNG2675C | 3 |

**Table 4:** The list of alternative transcripts in *H. salinarum* NRC-1.

| Chromosome | Start | End | Strand | Gene | Cluster |
| --- | --- | --- | --- | --- | --- |
| chr | 71317 | 71748 | forward | VNG0080H | 2 |
| chr | 716669 | 718771 | forward | VNG0940Gm | 3 |
| chr | 813372 | 814280 | forward | VNG1066C | 2 |
| chr | 1582844 | 1584007 | forward | VNG2149H | 4 |
| chr | 45456 | 44245 | reverse | VNG0051G | 2 |
| chr | 348898 | 348053 | reverse | VNG0452G | 2 |
| chr | 377209 | 376820 | reverse | VNG0485H | 2 |
| chr | 437350 | 436883 | reverse | VNG0564H | 2 |
| chr | 471549 | 469912 | reverse | VNG0615C | 3 |
| chr | 1011350 | 1010871 | reverse | VNG1355H | 2 |
| chr | 1315375 | 1314713 | reverse | VNG1775C | 3 |
| chr | 1502665 | 1502096 | reverse | VNG2036G | 3 |
| chr | 1634779 | 1633922 | reverse | VNG2204H | 2 |
| chr | 1731404 | 1730838 | reverse | VNG2321G | 2 |

**Table 5.**
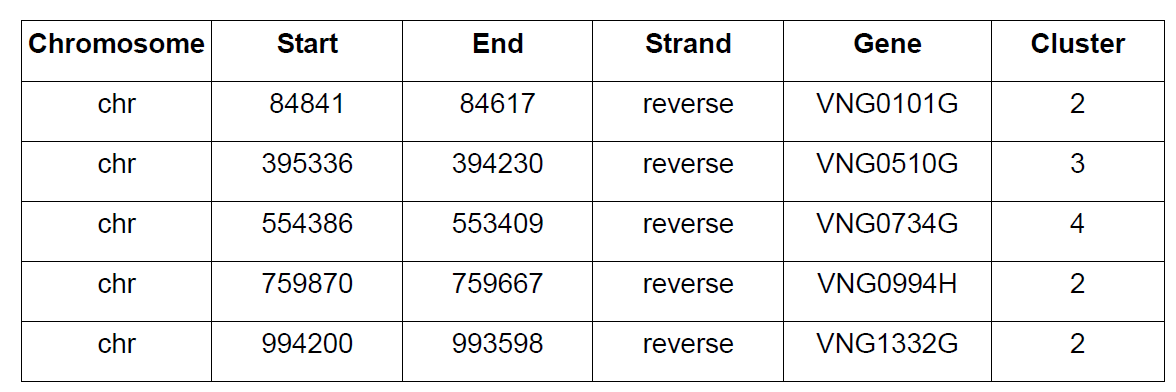
The list of Mis-annotation in *H. salinarum* NRC-1.

## Conclusions

In this work, we presented the intragenic differential expression in archaea by using computational techniques. We have several conclusions, which are as follows:

a). If you have a gene showing intragenic differential expression, but you do not know, you can design a probe in one of the half’s are think that this intragenic differential expression is valid for whole gene but it is not.
b). The supper GGB is useful for several additional things: the study of anti-sense RNAs. c).
c). Intragenic differential expression exists in archaea and is not just only for sotRNA, however also normal genes can have this. There are a lot of genes have intragenic differential expression and there are more than just sotRNA. In [6] presented a specific kind of transposes in just two conditions. In this work, we generalized this to all transposes families in several conditions.
d). A tool to spot mis-annotation of genome, for example the gene VNG0719G.
e). In [7] presented the TSSaRNAs and in this work, we also generalized the TSSaRNAs in archaea.

## Materials and Methods

To study and investigate the intragenic differential expression in archaea, we examined and analyzed all the publicly available gene expression data of archaea: *H. salinarum* NRC-1 at different condition i.e., growth curve, tiling arrays (GSE12923), *H. salinarum* NRC-1 vs TFB knockouts and synthetic TFB constructs (GSE31308), RNA expression data from *Halobacterium* NRC-1 in varied extracellular salinity conditions (GSE53544), *H. salinarum* NRC-1 vs VNG2099C knockout (GSE45988), evolution of context dependent regulation by expansion of feast/famine regulatory proteins [expression] (GSE61975), *Sulfolobus solfataricus P2*: growth curve, tiling arrays (GSE26779), *Pyrococcus furiosus* DSM 3638: growth curve, tiling arrays (GSE26782), *Methanococcus maripaludis* s2: growth curve, tiling arrays (GSE26777) [16]. In our analysis, we used all the above data, which we downloaded from the public databases and the datasets, which are not available in databases were collected from publications directly, to investigate the intragenic differential expression. We tabulated a brief description for each dataset in the supplementary Table 1.

The SOM and Gap statistics were used to report the intragenic differential expression in third domain of life archaea. Our method is defined as follows:

Step 1:

We took all the probes of a specific genes for each experiment, i.e., for each dataset, we have several experiments at different time.

Step 2:

In step 2, we used a technique of Gap statistics to estimate the number of clusters for each gene. From this step, we select only those genes in our analysis for next step, which have more than one clusters estimated by GAP statistics. In next step, we used SOM to clusters the probes for each gene to see the expression level.

Step 3:

In this step, we used SOM to clusters all probes of the gene. From this, we can see that some part of gene has over-expression and some part has lower-expression. We **didn’t used** RNA-seq data in our analysis to study the intragenic differential expression in archaea, however, we may clearly observe that if the tilling array for a gene shows over or lower-expressed, at the same position for RNA-seq data, also we can see that there is something important occur in same position (i.e., the signal breaks, etc).

Step 4:

We repeated our method for all genes of the third domain of life archaea to find all the genes, which have more than one clusters.

Step 5:

In this step, we did Eigen similarity search (BLAST search https://blast.ncbi.nlm.nih.gov/Blast.cgi) to eliminate the noise from our results. Since we have several genes which have more than one clusters i.e., the expression of transcripts in some part over-expressed and some part lower-expressed in other part, however, it maybe occurs due to Eigen similarity: one probe measures the expression in several positions. So therefore we did the Eigen similarity analysis to eliminate this noise, detail of this analysis in below section.

## Transcriptome analysis and re-normalization

We used all the publicly available data to present the intragenic differential expression in archaea. We downloaded the tilling microarray data of *H. salinarum* NRC-1 from NCBI and the GEO accession numbers are: GSE12923 [17], GSE31308 [18], GSE15788 [17], GSE45988 [19], GSE53544 [20] and GSE 61975 [21]. We re-normalized all the available data of *H. salinarum* NRC-1 up to date and visualized the re-normalized data at probe level by uploaded to GGB [15]. The *H. salinarum* NRC-1 growth curve (GSE12923) was normalized by comparing the Halobacterium NRC-1 reference sample with the experiment sample (growth curve) in [17], we re-normalized it with experiment sample t(0) sample. The *H. salinarum* NRC-1 vs TFB knockouts and synthetic TFB constructs (GSE31308) was normalized by comparing with reference sample. We re-normalized it with the t(0) experiment sample to t(1) experiment sample and so on. We compare our new re-normalized data to previous normalized published data, we found that our new re-moralized datasets MA-plot visualizations are much better than the previous published normalized data. Next we did the analysis for all available data to date by using computational techniques to investigate the intragenic differential expression in archaea.

We did normalization in a new way, as usual, normalization is done reference with experiments, however, we did analysis in way that make sense i.e., experiments with experiments in different way, the details are given in supplementary Table 2.

## Eigen-similarity and sequence similarity search in public databases

The Eigen similarity analysis was presented to split noise (artifacts) from the real biological results. We found that some genes have two or more clusters of *H. salinarum* NRC-1, however by Eigen match, we found that it may be due to noise Table 1, since some of the probes of that genes match in the genome more than once. We used a Bioconductor package to did the Eigen match analysis.

## Most significant genes

We select most important results i.e., the genes that make sense, from our data by using the Euclidean distance between clusters, which is defined as follows:

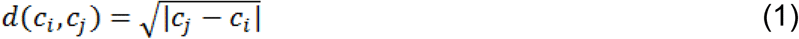

where c_*i*_ and c_*j*_ are number of clusters mean value in data. We define some threshold values to separate the genes in different groups. We have several genes that have two or more clusters, however we selected those genes which are more clear by using the above criteria. From this technique, we only selected those genes, which have clear two or more clusters, i.e., some part of genes clearly over-expressed and the other parts are clearly lower-expressed for detail see Figure 1 (a). The remaining genes, which have two or more clusters, however, which is not clear, we will consider them in our future work for further investigation.

## Availability of supporting data

The data sets supporting the results of this article are included within the article (and its additional file(s)) and third party repositories.

## Author Contributions

RZNV and AK analyzed data, interpreted data and wrote the manuscript. All authors discussed the biological findings and read/approved the final version of the present manuscript.

## Acknowledgments

Thanks to Dr Tie Koide for helpful discussions of the work. We also thank the Vencio and Tie labs members for helpful comments and feedback on this work. This work was supported by Projeto Jovem Pesquisador em Centros Emergentes da Fundação de Amparo à Pesquisa do Estado de São Paulo (FAPESP, http://fapesp.br/en/) [09/09532-0 to TK]; Edital Universal do Conselho Nacional de Desenvolvimento Científico e Tecnológico (CNPq) [473660/2013-0 to TK, 470120/2009-6 to TK, 476724/2013-9 to RZNV]; Fundação de Apoio ao Ensino, Pesquisa e Assistência do Hospital das Clínicas da Faculdade de Medicina de Ribeirão Preto da Universidade de São Paulo (FAEPA) [1640/2009 to TK]; Núcleo de Pesquisa em Ciência Genômica (NAP-CG) da Universidade de São Paulo; and fellowship FAPESP [2012/23329-5 to AK].

